# The copper chaperone ATOX1 exhibits differential protein-protein interactions and contributes to skeletal myoblast differentiation

**DOI:** 10.1101/2025.07.08.663731

**Authors:** Nathan Ferguson, Yu Zhang, Alexandra M. Perez, Allison T. Mezzell, Vinit C. Shanbhag, Michael J. Petris, Katherine E. Vest

## Abstract

Copper is an essential but potentially toxic nutrient required for a variety of biological functions. Mammalian cells use a complex network of copper transporters and metallochaperones to maintain copper homeostasis. Previous work investigating the role of copper in various disease states has highlighted the importance of copper transporters and metallochaperones. However, questions remain about how copper distribution changes under dynamic conditions like tissue differentiation. We previously reported that the copper exporter ATP7A is required for skeletal myoblast differentiation and that its expression changes in a differentiation dependent manner. Here, we sought to further understand the ATP7A-mediated copper export pathway by examining ATOX1, the copper chaperone that delivers copper to ATP7A. To investigate the role of ATOX1 in a dynamic cellular context, we characterized its binding partners during myoblast differentiation using the proximity labeling protein APEX2 to biotinylate proteins near ATOX1. We discovered that the ATOX1 interactome undergoes dramatic changes as myoblasts differentiate. These dynamic interactions correlate with distinct phenotypes of ATOX1 deficiency in proliferating and differentiated cells. Together, our results highlight the dynamic interactome of ATOX1 and its contribution to myoblast differentiation.

## Introduction

Copper (Cu) is a vital trace nutrient required for a variety of biological processes. It serves as a catalytic cofactor in cytochrome c oxidase during cellular respiration, supports antioxidant defense via superoxide dismutase (SOD1), facilitates extracellular matrix remodeling through lysyl oxidases (LOX), and modulates cellular signaling by allosteric regulation of kinases such as MEK1/2.^1^ However, unregulated copper levels can be toxic. Excess copper can induce oxidative stress, disrupt the function of other metalloproteins, or trigger cell death through a recently discovered pathway known as cuproptosis.^2,3^ To maintain copper homeostasis, cells rely on a network of transporters and chaperones that tightly control import, export, and intracellular distribution. In mammalian cells, the primary importer is CTR1, which facilitates the uptake of Cu^+1^ ions which are then transferred to various metallochaperones including ATOX1.^4^ The ATOX1 protein delivers copper to the primary exporter proteins, ATP7A and ATP7B.^5^ ATP7A is expressed in most non-hepatic tissues and is primarily localized to the *trans*- Golgi network where it provides copper to secreted cuproproteins.^1^ Under high copper conditions, ATP7A and ATP7B translocate to the cell membrane to move intracellular copper directly to the extracellular space.^6,7^ Thus, in mammals, ATP7A and ATP7B are critically important for both copper mobilization and detoxification.

Disruptions in copper homeostasis can give rise to serious disease. Loss-of-function mutations in *ATP7A* lead to Menkes disease (MD), a fatal, infantile-onset copper deficiency characterized by hypotonia, connective tissue defects, neurodegeneration, and failure to thrive.^8^ The severity of Menkes disease symptoms and overall survival is known to vary, even between related individuals with the same mutations in *ATP7A*. Given the pleiotropic nature of symptoms arising from Menkes and other copper-related diseases, understanding dynamic cellular copper metabolism across multiple tissue types and differentiation states is of utmost importance.

Research from various groups has shown that copper is important for early brain development, contributing to myelination, synaptic connectivity, and neurotransmitter synthesis.^9,10,11^ Consistent with the important role of copper in brain development, early supplementation in infants with Menkes disease has been shown to partially rescue developmental delays.^8^ We and others have also shown that copper is important for differentiation of skeletal muscle, osteoblasts, and hematopoietic progenitor cells.^12,13,14,15,16,17^ Collectively, these studies highlight the essential and multifaceted roles copper plays across multiple cell and tissue types during differentiation and development. However, many questions remain surrounding the mechanism by which copper distribution is regulated within different cellular environments, particularly under shifting metabolic conditions.

To address these questions, our group uses skeletal muscle as a model system due to its well-characterized differentiation pathway. Skeletal muscle contains about 23% of total body copper and is notably affected in Menkes disease, where severe muscle weakness is a hallmark symptom.^1,8,18^ In previous work, we explored the molecular requirements of copper during skeletal muscle differentiation and found that the copper dependent enzyme lysyl oxidase (LOX), which is required for extracellular matrix development, plays a vital role in the differentiation of skeletal myoblasts to mature myotubes.^12,13^ Notably, ATP7A is required for myotube formation as it provides copper to LOX, and thus deficiency in copper, ATP7A, or LOX activity impairs myotube formation. The present study focuses on the role of ATOX1, the primary copper chaperone for ATP7A, during muscle differentiation.^19,20^ ATOX1 and its homologs were originally identified as antioxidant proteins, with many studies demonstrating that ATOX 1 expression increases in response to oxidative stress, mitigating cellular damage.^10,21^ More recently, ATOX1 has been shown to indirectly promote copper dependent activation of the MEK/ERK signaling cascade in human BRAF^V600E^ expressing cancer cells, while the copper chaperone for SOD1 (CCS) is responsible for direct copper delivery to MEK1/2.^22,23^ Other studies have implicated ATOX1 as a putative transcription factor that controls the expression of the cyclin-D pathway.^24,25^ These diverse roles highlight the functional importance of ATOX1 across multiple cell types. However, significant gaps remain in our understanding of how ATOX1 function varies across cellular contexts, particularly during dynamic processes such as differentiation.

In this study, we investigate the multifaceted role of ATOX1 during skeletal muscle differentiation. To do so, we generated stable skeletal myoblast lines that express a construct containing the proximity labeling protein APEX2 fused to the amino (N)-terminus of ATOX1 (APEX2-ATOX1) under the control of an inducible promoter. This construct differs from the previously studied carboxy (C)-terminal tagged ATOX1 which contained both APEX2 and an exogenous nuclear localization signal (NLS).^26^ Using comparative proteomics across different stages of myoblast differentiation, we identified dynamic changes in APEX2-ATOX1 proximal proteins. Additionally, we found that loss of ATOX1 leads to increased myoblast proliferation and a cell density-dependent defect in differentiation. These fundings suggest that ATOX1 functions are differentiation state-dependent in skeletal muscle cells and support the broader model that ATOX1 mediates diverse functions across mammalian tissues.

## Results

### APEX2-ATOX1 construct production and expression

To elucidate the functional importance of ATOX1 in skeletal myoblast differentiation, we sought to identify its proximal protein binding partners using a proximity labeling approach. The APEX2 protein is an engineered ascorbate peroxidase that biotinylates proteins within approximately 20 nm of the bait protein.^27^ A construct encoding a DYKDDDDK (Flag) tag, APEX2, and a Myc tag at the amino (N) terminal end of human ATOX1 was expressed under the control of a CMV promoter in pcDNA 3.1 (Figure 1A). Expression of the construct was confirmed by immunoblot with antibodies targeting the Flag tag and was further validated using antibodies targeting human and mouse ATOX1, which detected a ∼40 kDa band corresponding to the fusion protein, in contrast to the ∼7 kDa endogenous mouse Atox1 (Figure 1B). Optimal labeling conditions were determined through titrations of biotin-phenol (BP) and hydrogen peroxide (H2O2) across various time points. Immunoblot analysis using streptavidin-HRP to identify biotinylated proteins revealed that 2.5mM BP, 0.75mM H2O2, and 60 seconds time were optimal for robust labeling (Figure 1C-E).

**Figure 1:**
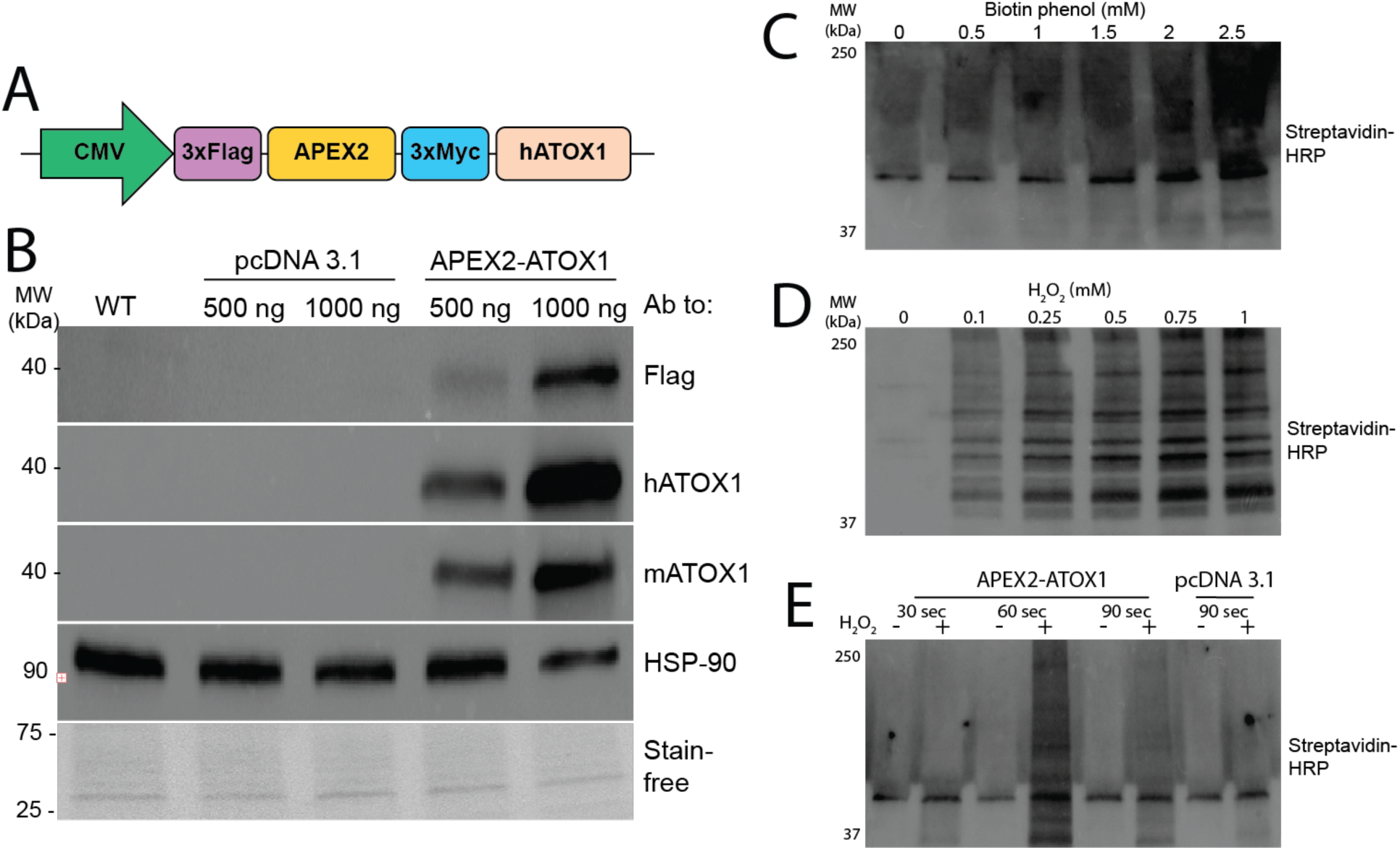
Generation of APEX2-ATOX1 constructs ***A)*** The APEX2-ATOX1 construct contains 3x Flag and 3x Myc tags flanking the full-length APEX2 sequence at the amino terminal of the human ATOX1 protein. All tags are separated by glycine-serine hinges. Constructs were subcloned into a pcDNA 3.1 plasmid under the control of the cytomegalovirus immediate early promoter (CMV). ***B)*** Immunoblot of lysates from C2C12 myoblasts transfected with APEX2-ATOX1constructs showing detection of APEX2-ATOX1 fusion constructs by antibodies to Flag, human (h) or mouse (m) ATOX1. Antibodies to HSP-90 and total proteins detected using stain-free gel imaging technology (Bio-Rad) were used as loading controls. Labeling efficiency was determined by probing blots with horseradish peroxidase (HRP) conjugated streptavidin in cells titrated with biotin phenol (***C***), hydrogen peroxide (***D***), or at variable time points (***E***).

To eliminate variability from uneven transfection efficiency of pcDNA 3.1 plasmids in C2C12 cells, we generated stable C2C12 myoblast lines expressing APEX2-ATOX1 under the control of a doxycycline-inducible TRE promoter (Figure 2A). Cells were transduced and selected with puromycin to produce stable cells and doxycycline titration confirmed dose-dependent induction of APEX2-ATOX1 expression (Figure 2B). A doxycycline concentration of 2.5 µg/mL was optimal for robust APEX2-ATOX1 expression in myoblasts, myocytes, and myotubes (Figure 2 B-D). Although N-terminal epitope tagging has been used to study ATOX1, we validated that the APEX2 tag does not alter its subcellular localization.^28^ Fractionation followed by immunoblot analysis revealed that endogenous ATOX1 and APEX2-ATOX1 localize primarily to the cytosol in myoblasts (Figure 2E), myocytes (Figure 2F), and myotubes (Figure 2G). Although trace amounts of APEX2-ATOX1 were detected in nuclear fractions, endogenous ATOX1 was not, though low abundance ATOX1 may fall below the detection threshold of immunoblotting (Figure 2 E-G). These localization patterns align with previous observations in neuronal differentiation, where endogenous ATOX2 remained cytosolic throughout the.^10^ Collectively, these results confirm that the APEX2-ATOX1 fusion construct mimics the localization of endogenous ATOX1.

**Figure 2:**
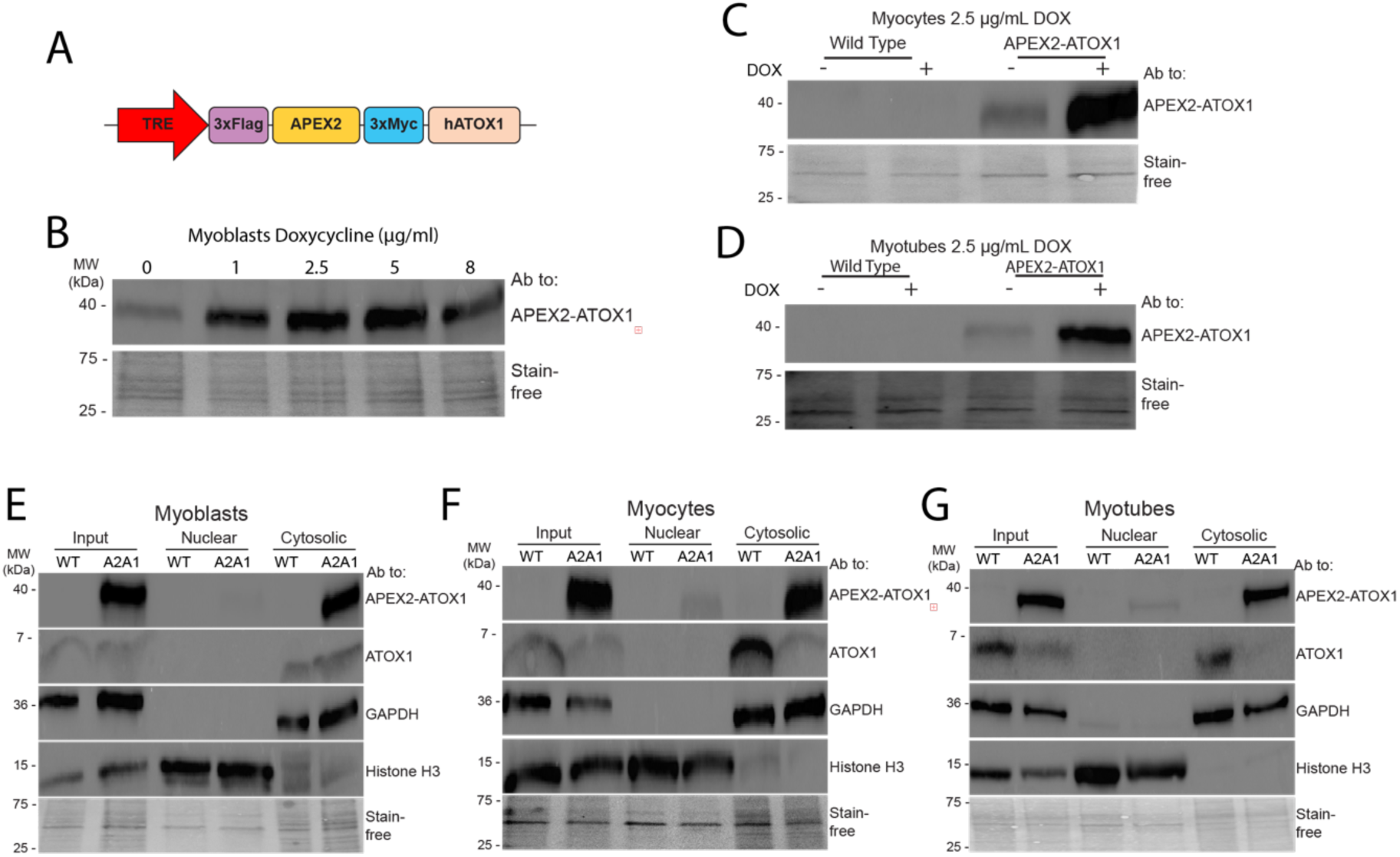
Stable cells express inducible APEX2-ATOX1 ***A*)** Stable C2C12 cells were generated that express the APEX2-ATOX1 fusion construct under the control of a tetracycline responsive element (TRE). ***B*)** Doxycycline (DOX) titration revealed the optimal concentration to be 2.5 µg/ml as determined by immunoblot using an antibody to ATOX1. DOX at 2.5 µg/ml was used to indue APEX2-ATOX1 expression in myocytes (***C***) and myotubes (***D***). Localization of APEX2-ATOX1 construct (A2A1) as determined by fractionation in myoblasts (***E***), myocytes (***F***), and myotubes (***G***) as analyzed by immunoblot using antibodies to GAPDH to indicate the cytosolic fraction and histone H3 to indicate the nuclear fraction. Total protein as detected by stain-free gel imaging technology (Bio-Rad) was used as a loading control.

### ATOX1 interacts with canonical and noncanonical binding partners in a differentiation dependent manner

To characterize the ATOX1 interactome, we performed comparative proteomics with label-free quantification to identify proteins in proximity to APEX2-ATOX1. Construct expression was induced in myoblasts (MB), myocytes (MC), and myotubes (MT), and biotinylated proteins were isolated from lysates by streptavidin-based capture in three independent experiments per differentiation state (Figure 3A). Induced cells without biotin phenol treatment were used as negative controls. Protein peaks were filtered to include only those with high confidence (false discovery rate of 99%) and proteins identified in -BP controls were filtered out yielding 708 total proteins (Supplemental Table 2). Proteins that were detected in all three replicates (and not detected in -BP controls) for each differentiation state were included for downstream functional annotation yielding 212, 99, and 76 ATOX1 proximal proteins in myoblasts, myocytes, and myotubes (Figure 3B). Functional annotation of ATOX1 proximal proteins was performed using the NIH database for annotation, visualization, and integrated discovery (DAVID).^29^ Top GO terms for biological processes and molecular function and top KEGG pathways varied with differentiation state (Figure 3C-E). In myoblasts, ATOX1 proximal proteins are enriched in pathways related to cell cycle regulation and insulin signaling, reflecting a proliferative profile (Figure 3C). In myocytes, enrichment was observed in categories related to endomembrane trafficking and protein binding/bridging were enriched, which is consistent with the role of ATOX1 in delivering Cu to ATP7A (Figure 3D). In myotubes, ATOX1 proximal proteins are primarily enriched in functions related to mRNA binding, processing, and transport (Figure 3E). Comparable enrichment patters were also observed using the STRING database for pathway analysis (Supplemental Figure 1).^30^ Collectively, these findings suggest that the ATOX1 protein interaction network dynamically remodels in response to cellular differentiation state.

**Figure 3:**
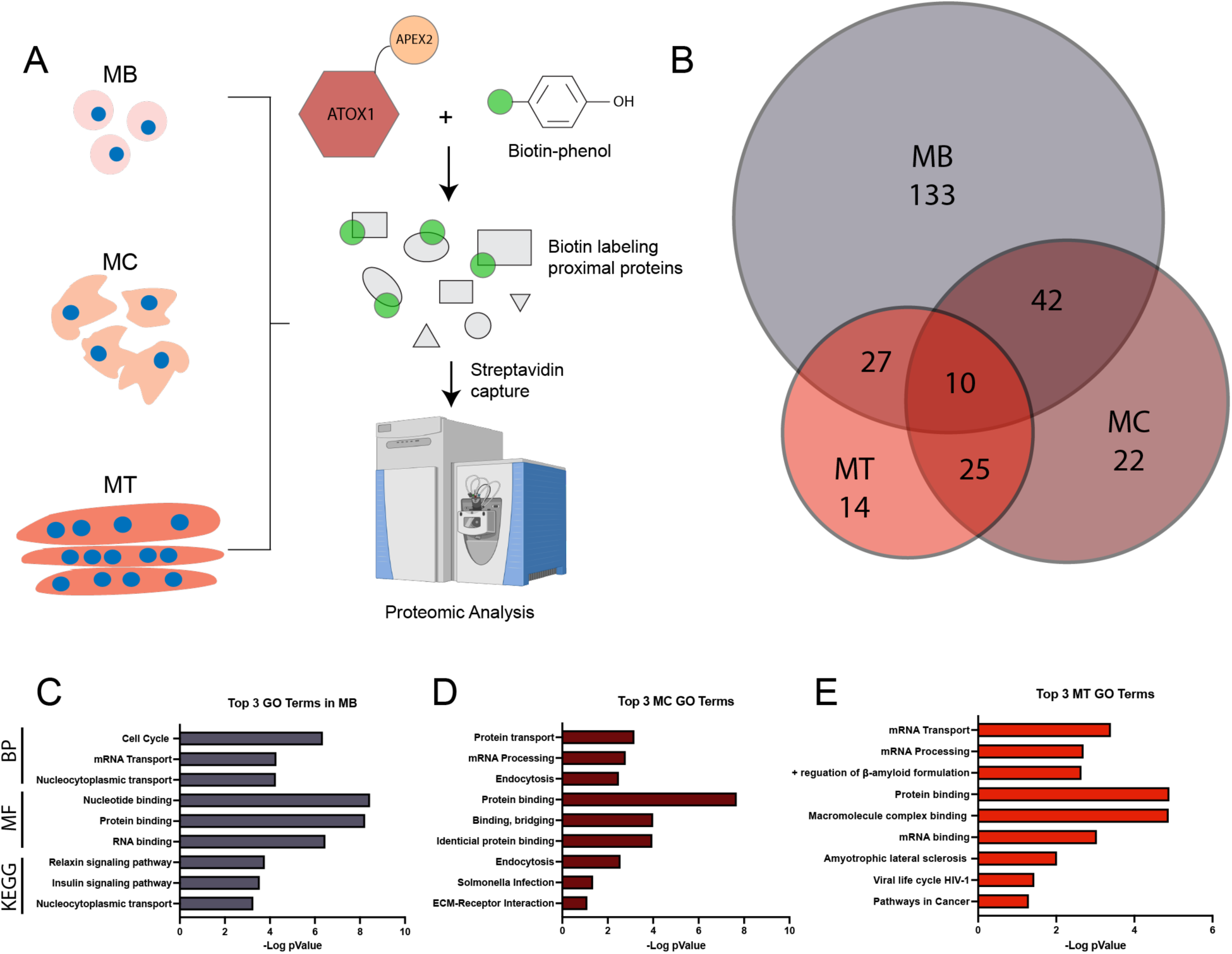
Comparative proteomics and label-free quantification reveals differential ATOX1 proximal proteins detected during myoblast differentiation. ***A)*** Expression of APEX2-ATOX1 was induced in myoblasts (MB), myocytes (MC), and myotubes (MT). After labeling, lysates were isolated with streptavidin beads and analyzed by comparative proteomics. Cells without biotin phenol were used as negative controls and three independent experiments were performed per differentiation condition. ***B)*** Venn diagram of proteins detected in all three replicates of MB, MC, and MT that were not detected in negative control cells without biotin phenol. These hits were used for gene ontology and pathway analysis. The top three hits each for GO_Biological Process (BP), GO_Molecular Function (MF), and the Kyoto Encyclopedia of Genes and Genome (KEGG) pathways are shown for MB (***C***), MC (***D***), and MT (***E***).

Both known and putative ATOX1 interacting proteins were detected in APEX2-ATOX1 proximity labeling data sets. Recent studies identified cysteine rich intestinal protein 2 (CRIP2) as a copper-binding transcription factor that interacts with ATOX1 and plays a role in copper dependent skeletal myoblast differentiation.^26,31^ CRIP2 was detected as an ATOX1 proximal protein in proteomic data (Supplemental Table 2) and validated by immunoblotting independent streptavidin eluates from myoblasts and myocytes (Figure 4A, B). Several proteins detected in the previous ATOX1-APEX2-NLS study that identified CRIP2 were also detected as ATOX1 proximal proteins in this study including the nucleic acid binding proteins FUBP1, HNRNPAB, and SYNCRIP/HNRNPQ (Supplemental Table 2).^26^ Immunoblot analysis of streptavidin eluates revealed that SYNCRIP was most easily detected at the myocyte stage, though enrichment over minus biotin phenol negative control was not significant (Figure 4A, C). Interestingly, though no studies have shown a direct interaction between ATOX1 and MEK1/2 kinases, they have been functionally linked in the context of copper dependent MEK-ERK signaling.^22^ Both MEK1 and MEK2 were detected as APEX2-ATOX1 proximal proteins in myoblasts in comparative proteomics (Supplemental Table 2) and confirmed by immunoblot (Figure 4A, D), suggesting that MEK1/2 kinases may bind ATOX1 in muscle cells. Some known ATOX1 binding proteins were not detected in the APEX2-ATOX1 proximal proteome data including the copper chaperone for SOD1 (CCS), which can bind and undergo copper transfer with ATOX1.^32^ CCS interaction with APEX2-ATOX1 was however detected by immunoblot in myoblasts and myocytes (Figure 4A, E). Similarly, ATP7A was not detected in proteomics data, which is likely due to the use of boiling during elution from streptavidin beads, which is known to cause ATP7A aggregation.

**Figure 4:**
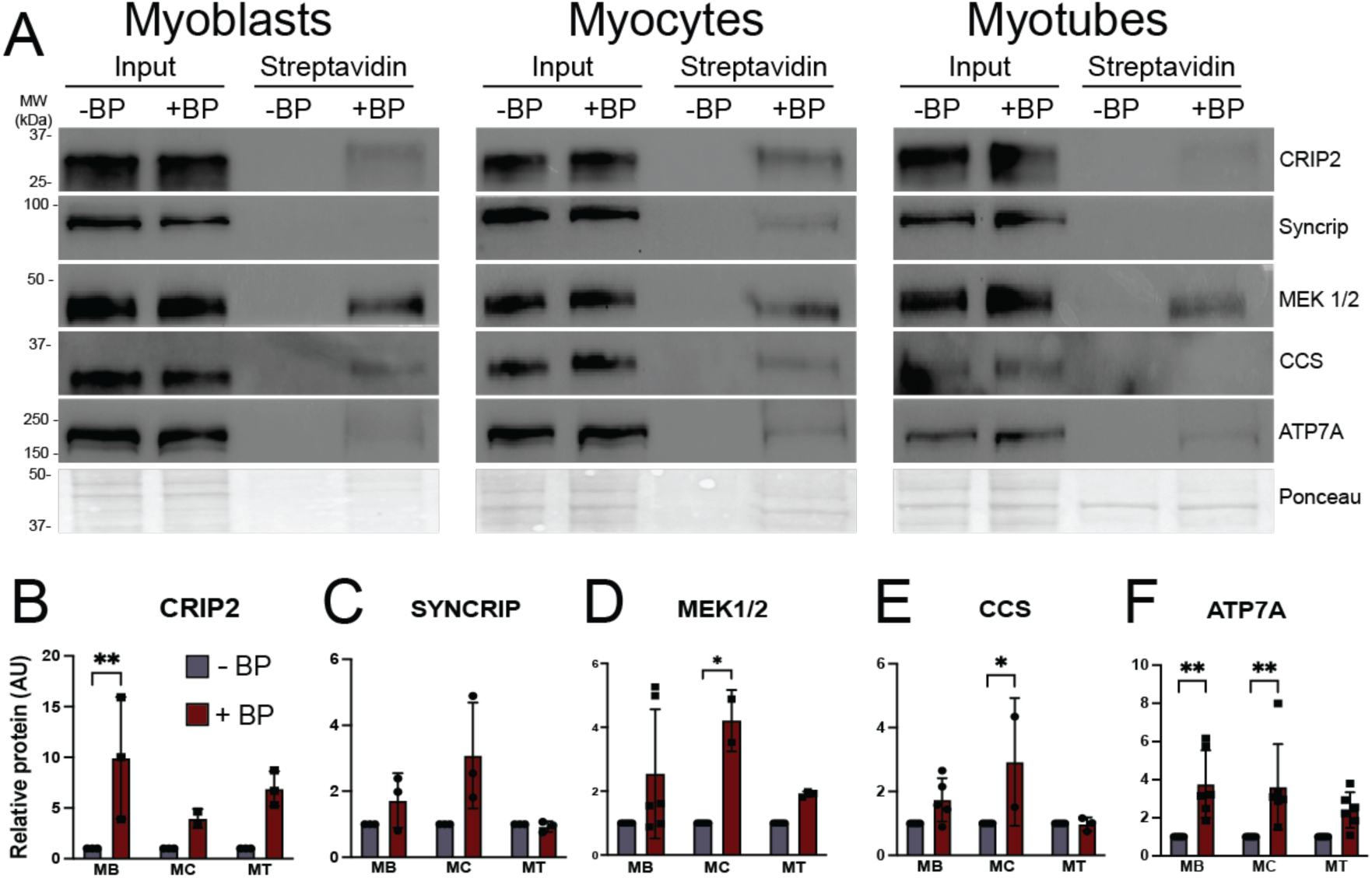
Shifting ATOX1 binding partners during myoblast differentiation as detected by immunoblot ***A)*** Representative blots showing streptavidin bead elutions from myoblasts (MB), myocytes (MC), and myotubes (MT) expressing APEX2-ATOX1 and treated with biotin phenol (BP). Cells without BP (-BP) were used as negative controls. Total protein as detected by Ponceau stain was used as loading control. ***B-F)*** Densitometric quantification of candidate ATOX1 binding proteins from blots shown in (***A***). Shown for all is mean ± standard deviation for n = 2-5 experiments. Statistical significance was determine using two-way ANOVA with Sidak post-hoc correction for multiple comparisons. * p < 0.05, ** p < 0.01

However, elution using urea containing buffer without boiling allowed for detection of significantly enriched ATP7A in myoblasts and myocytes (Figure 4A, F). ATP7A was also detectable in myotubes (Figure 4A), though enrichment over -BP control was not significant (Figure 4F). Taken together, these results suggest that while ATOX1 fulfills its well-established canonical functions in the intermediate stages of differentiation (myocytes), it may also participate in distinct regulatory processes in proliferating myoblasts and fully differentiated myotubes.

### ATOX1 deficiency impacts myoblast proliferation and differentiation

In myoblasts, functional annotation revealed enrichment for gene ontology terms related to cell cycle. In previous studies using mouse embryonic fibroblasts (MEFs) and BRAF^V600E^ mutant melanoma cell lines, copper was shown to promote proliferation by binding to and potentiating the activity of MEK1/2 kinases leading to increased phosphorylation of the downstream MEK targets ERK1/2 kinases. ATOX1 also promotes growth of BRAF^V600E^ expressing melanoma cells, though its effect on ERK1/2 phosphorylation was proposed to be indirect.^22^ In our study, both MEK1/2 kinases were detected as APEX2-ATOX1 proximal proteins in myoblasts (Figure 4A and Supplementary Table 2). However, neither siRNA-mediated ATOX1 deficiency nor APEX2-ATOX1 overexpression affected MEK1/2 kinase activity as assessed by ERK1/2 phosphorylation levels (Figure 5A, B). Similarly, immunoprecipitation of MEK1/2 kinases enriched for ERK1/2 kinases but did not yield any detectable ATOX1 protein, even in the presence of insulin stimulation of MEK/ERK signaling (Figure 5C). ATOX1 knockdown did, however, lead to a small but significant increase in myoblast growth as detected by cell counting (Figure 5 D, E). ATOX1 deficiency also correlated with a significant decrease in CCS levels (Figure 5D), suggesting increased copper availability.

**Figure 5:**
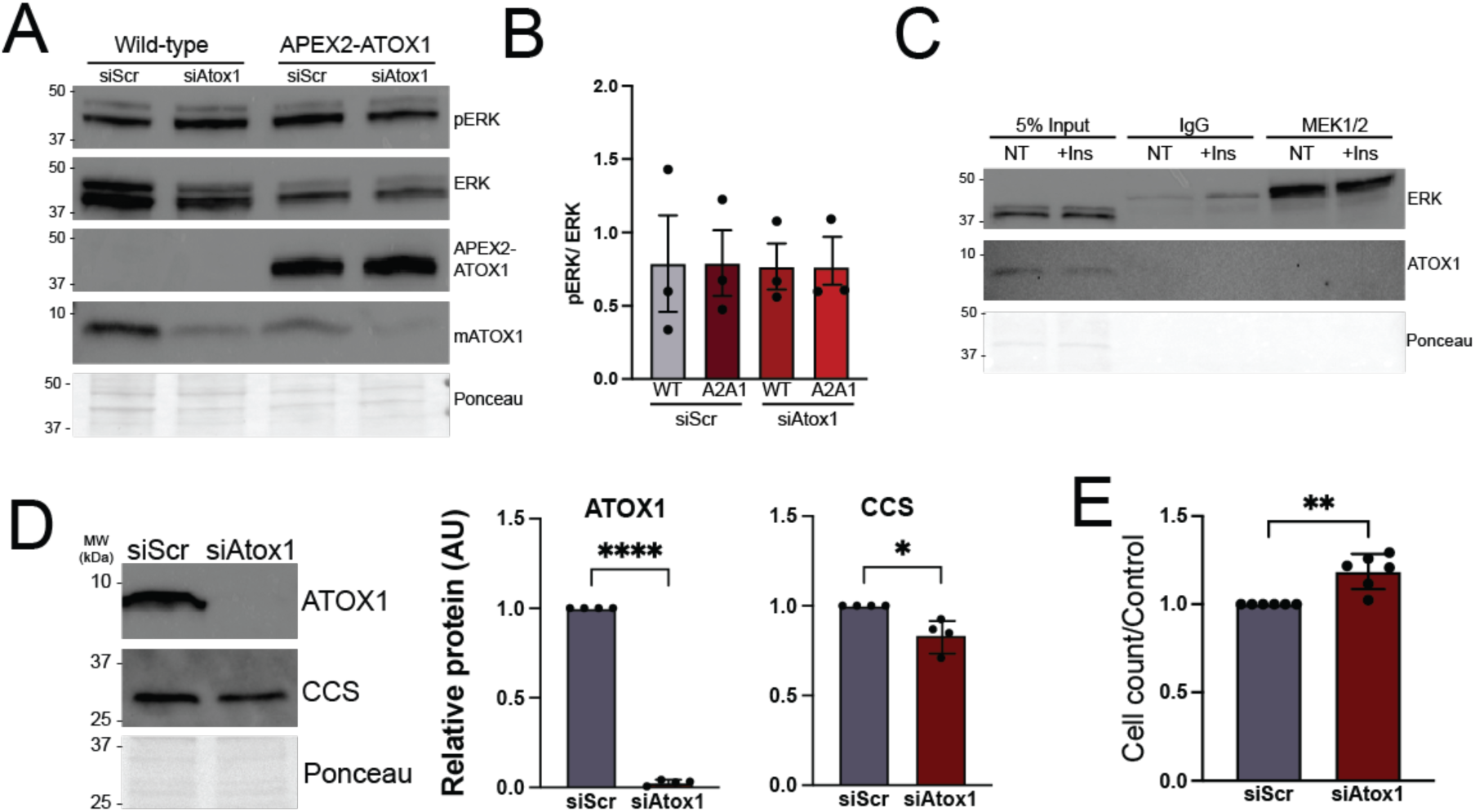
ATOX1 deficiency impacts myoblast proliferation but not ERK phosphorylation ***A)*** Representative immunoblot showing phosphorylated ERK1/2 (pERK) and total ERK1/2 (ERK) in wild type or APEX2-ATOX1 expressing myoblasts transfected with negative control siRNA (siScr) or *Atox1* targeting siRNA (siAtox1). Blots probed with an antibody to ATOX1 were used to show expression of APEX2-ATOX1 and endogenous ATOX1 knockdown. Total protein as detected by Ponceau stain was used as a loading control. ***B)*** Ratio of pERK/ERK in wild type (WT) and APEX2-ATOX1 (A2A1) expressing myoblasts with (siAtox1) and without (siScr) *Atox1* knockdown. Shown is mean ± standard deviation for quantification of n = 3 immunoblots. ***C)*** Representative immunoblot of immunoprecipitation (IP) using an antibody to MEK1/2 in myoblasts with (+Ins) or without (NT) insulin treatment to stimulate MEK/ERK signaling. The IP was probed with an antibody to ERK1/2 (ERK) showing interaction between MEK1/2 and ERK1/2 and an antibody to ATOX1 showing no interaction. Total protein as detected by Ponceau stain was used as a loading control. ***D)*** Representative immunoblot probed with an antibody to ATOX1 to show ATOX1 knockdown in myoblasts and an antibody to CCS in control (siScr) and *Atox1* knockdown (siAtox1) myoblasts. Quantifications show mean ± standard deviation for n = 4 experiments. Statistical significance was determined by t-test. ***E)*** Proliferation as determined by cell counting in control (siScr) and *Atox1* knockdown (siAtox1) myoblasts. Shown is mean ± standard deviation for n = 6 experiments. Statistical significance was determined by t-test. * p < 0.05, ** p < 0.01. **** p < 0.0001.

Our group previously demonstrated that ATP7A levels increase dramatically during the early stages of myoblast differentiation before declining as cells mature into fully differentiated myotubes and that ATP7A deficiency severely impairs myotube formation.^12^ Unlike ATP7A, levels of ATOX1 protein (Figure 6A, B) and RNA (Figure 6C) increase gradually throughout myoblast differentiation and remain high.^12^ Previous studies of ATOX1 revealed that its ability to bind copper is regulated by intracellular redox balance, which shifts ATOX1 to a more reduced state during neuronal differentiation.^10^ Thiol labeling by EZ-link maleimide-PEG11-biotin allows for detection of free reduced or oxidized/bound thiols in ATOX1 by tracking the shift in molecular weight (∼1kDa) after labeling by maleimide-PEG11-biotin. There are three cysteine residues in ATOX1, two of which contribute to the copper-bridged homodimerization and form a disulfide bond in an oxidizing environment, while the third cysteine is expected to remain free.

**Figure 6:**
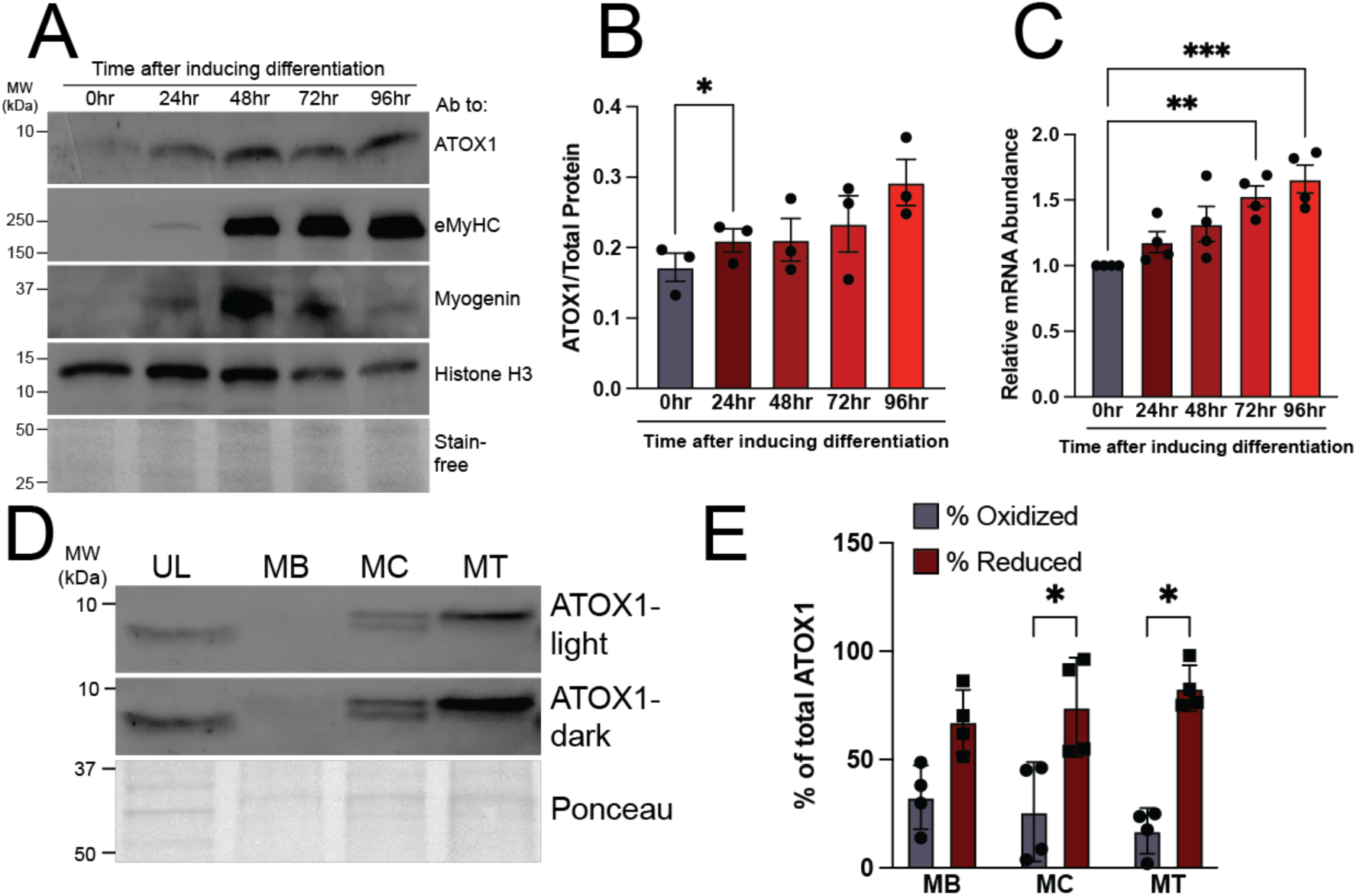
ATOX1 levels increase and redox state fluctuates during myoblast differentiation ***A)*** Representative immunoblot showing increasing levels of ATOX1 at 24, 48, 72, and 96 hours after inducing differentiation corresponding to the myocyte (24-48), myocyte/early myotube (48- 72), and mature myotube (72-96) stages. Myogenin and embryonic myosin heavy chain (eMyHC) used as intermediate and late-stage differentiation markers, respectively. Histone H3 and total protein as detected by stain-free gel imaging technology (Bio-Rad) used as loading controls. ***B)*** Quantification showing increased ATOX1 relative to total protein (stain-free) during differentiation. Shown is mean ± standard deviation of immunoblot quantification for n = 3 experiments. Statistical significance was determined by one-way ANOVA with Dunnett post-hoc test for multiple comparisons. * p < 0.05. ***C)*** qRT-PCR data showing increased levels of *Atox1* transcript during differentiation. Shown is mean ± standard deviation for n = 4 experiments. Statistical significance was determined by one-way ANOVA with Dunnett post-hoc test for multiple comparisons. ** p < 0.01, *** p < 0.001. ***D)*** Representative immunoblot showing unlabeled (UL) and EZ-link maleimide PEG11-biotin labeled ATOX1 in myoblasts (MB), myocytes (MC), and myotubes (MT). Short (ATOX1-light) and long (ATOX1-dark) exposures are shown to highlight reduced (top band) and oxidized (bottom band) ATOX1 in MB, MC, and MT lanes. Total protein as detected by Ponceau stain is used as a loading control. ***E)*** Quantification of percent oxidized and reduced ATOX1 as calculated relative to total ATOX1 (top band + bottom band). Shown is mean ± standard deviation for n = 4 experiments. Statistical significance was determined by two-way ANOVA with Tukey’s post-hoc testing for multiple comparisons. * p < 0.05.

Both labeled and unlabeled ATOX1 could be detected at each stage of differentiation, although the results were highly variable (Figure 6D). Quantitative analysis revealed no significant change in labeled ATOX1 across myoblast differentiation, but significant enrichment for labeled (reduced) compared to unlabeled (oxidized/bound) ATOX1 was only detectable in myocytes and myotubes (Figure 6E). These results suggest that thiol oxidation in ATOX1 fluctuates in multiple differentiation states but may be enriched in more differentiated myocytes and myotubes.

As previously reported, ATOX1 deficiency a marked defect in myoblast differentiation (Figure 7A, B). However, myotubes were significantly thinner (Figure 7C) and levels of embryonic myosin heavy chain were reduced (Figure 7D) in ATOX1 deficient cells, suggesting that late stages of differentiation such as secondary fusion or hypertrophy may depend on ATOX1 activity.^12^ Given the established function of ATOX1 delivering copper to secreted cuproenzymes, we hypothesized that this relatively mild differentiation phenotype in ATOX1 deficient cells may be caused by a lack of secreted cuproproteins that would have a stronger phenotypic effect in low cell density conditions. To test this hypothesis, we induced differentiation in control and *Atox1* knockdown myoblasts plated at low density. Under these conditions, ATOX1 deficiency led to a significant impairment in myoblast differentiation as indicated by a reduced fusion index (Figure 7E, F), formation of smaller myotubes (Figure 7G), and a trend towards decreased embryonic myosin heavy chain levels (Figure 7H). Notably, no overt phenotype was detected when ATOX1 deficiency was induced in fully differentiated myotubes (Supplemental Figure 2). Taken together, these results support a model in which high cell density enables low levels of residual ATOX1 or ATOX1-independent copper delivery to ATP7A to sustain secreted cuproprotein activity and promote myotube formation.

**Figure 7:**
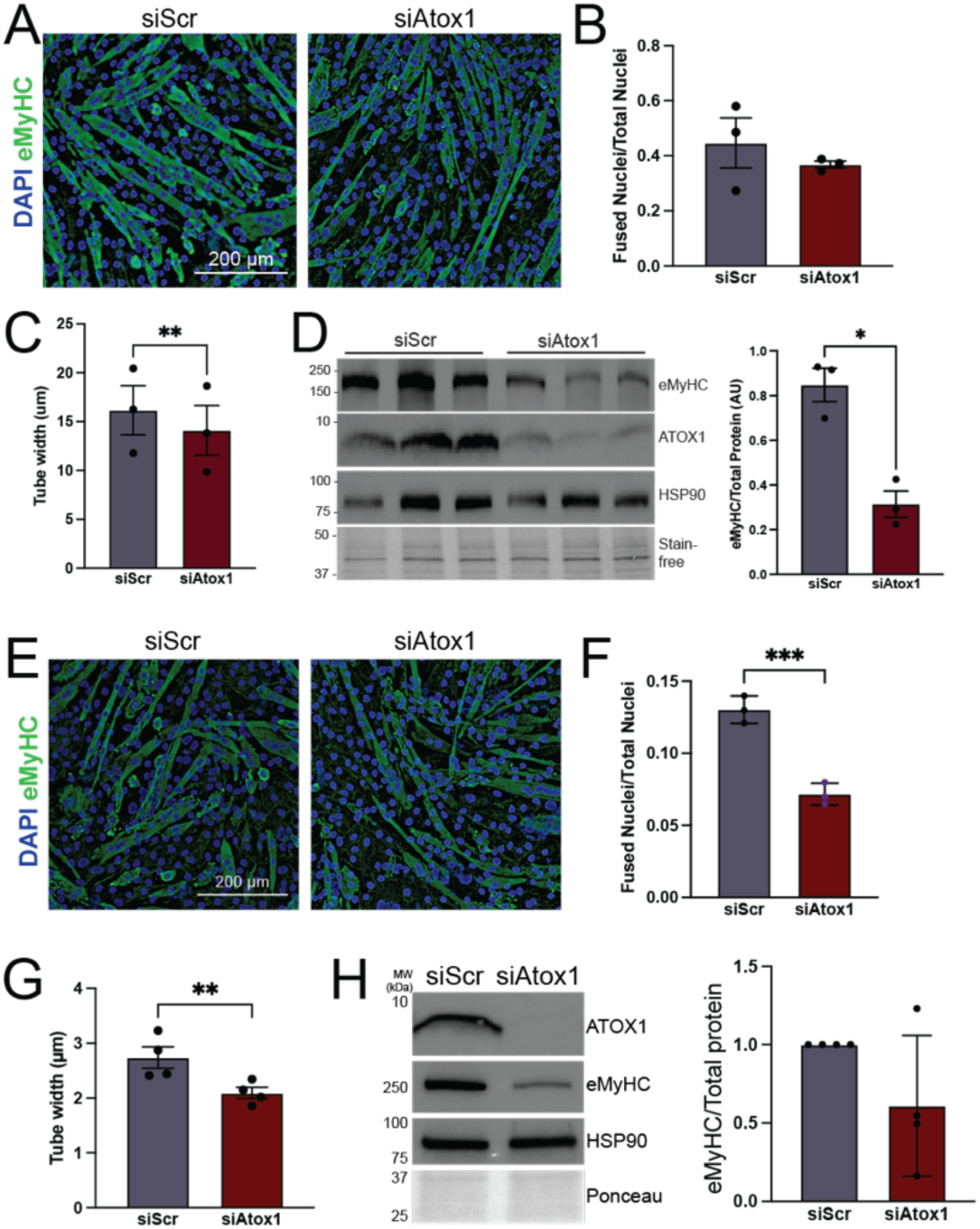
ATOX1 deficiency causes a cell density dependent differentiation defect ***A)*** C2C12 myotubes from control (siScr) or ATOX1 deficient (siAtox1) cells plated at high density. Myotubes are stained with an antibody to embryonic myosin heavy chain (eMyHC, green) and nuclei are stained with DAPI (blue). ***B)*** Fusion index (fraction of nuclei in myotubes relative to total nuclei) from high-density siScr and siAtox1 myotubes. ***C)*** Tube width measurement from high density siScr and siAtox1 myotubes. ***D)*** Immunoblot of lysates from high-density siScr and siAtox1 myotubes probed with antibodies to eMyHC as a measure for differentiation and ATOX1 to show knockdown. HSP-90 and total protein detected using stain-free gel imaging technology (Bio-Rad) were used as loading controls. Shown are lysates from three independent experiments. Quantification shows levels of eMyHC normalized to total protein. ***E)*** C2C12 myotubes from siScr and siAtox1 cells plated in low density and stained with an antibody to eMyHC and DAPI. ***F)*** Fusion index from siScr and siAtox1 myotubes plated at low density. ***G)*** Tube width from siScr and siAtox1 myotubes plated at low density. ***H)*** Representative immunoblot of lysates from high-density siScr and siAtox1 myotubes probed with antibodies to eMyHC as a measure for differentiation and ATOX1 to show knockdown. HSP-90 and total protein as detected by Ponceau stain were used as loading controls. Quantification shows levels of eMyHC normalized to total protein. For all, shown is mean ± standard deviation for n = 3-4 experiments. Statistical significance was determined by t-test. * p < 0.05, ** p < 0.01, *** p < 0.001.

## Discussion

In this study, we aimed to elucidate how copper is delivered to ATP7A during myoblast differentiation by characterizing the differentiation dependent protein binding partners and activities of ATOX1, the copper chaperone for ATP7A. To this end, we generated stable C2C12 cells expressing an inducible APEX2-ATOX1 proximity labeling construct and discovered that the ATOX1 interactome undergoes dramatic changes as differentiation progresses. These shifting protein-protein interactions correlate with variable ATOX1 functions, wherein ATOX1 appears to suppress myoblast proliferation while promoting myoblast differentiation in a cell density dependent manner. The results reported here agree with a growing list of studies suggesting that variable ATOX1 regulation and activities contribute to tissue differentiation, development, and cancer.^10,22,25,33,34^

In proliferating myoblasts, ontologies related to cell cycle and insulin signaling, which included MEK1/2 kinases, were enriched in ATOX1-proximal proteins. The MEK/ERK signaling pathway is active in proliferating myoblasts but must decrease to promote differentiation into myotubes.^35^ Although ATOX1 has been implicated in promoting copper dependent MEK1/2 activity, it was proposed to play an indirect role.^22^ By contrast, the copper chaperone for SOD (CCS) is the likely delivery pathway for copper to MEK1/2 and other kinases.^23^ Although an ATOX1-MEK1/2 interaction was suggested by proximity labeling, we were unable to detect ATOX1 by reverse-immunoprecipitation using an antibody to MEK1/2. However, few studies have reported successful immunoprecipitation of endogenous ATOX1, so a transient interaction between these proteins cannot be ruled out. In ATOX1 deficient myoblasts, no significant effect on MEK1/2 kinase activity was detected, though surprisingly, some of our data (Figure 5B and not shown) suggest a slight increase in phosphorylated ERK in *Atox1* knockdown cells. While this result could be related to an inhibitory function of ATOX1 on MEK1/2, we consider it is more likely related to increased copper availability in ATOX1 deficient cells. The reduced levels of CCS protein following *Atox1* knockdown are consistent with increased copper availability. ATOX1 proximal proteins in myoblasts also include negative regulators of the cell cycle like ERBB receptor feedback inhibitor 1 (ERRFI1) and transforming growth factor beta regulator 1 (TBRG1), both of which inhibit cancer cell growth.^36,37^ Thus, ATOX1 may play additional undefined roles in regulating proliferation of myoblasts and other cell types. Indeed, a role for ATOX1 in tumor growth and metastasis has been identified in breast and colon cancers.^22,34,38,39^ However, in HEK293T and MDA-MB-231 breast cancer cells investigators detected a substantial interaction between ATOX1 and components of the anaphase promoting complex, a key regulator of cell proliferation, that were not detected in this study.^40^ This discrepancy may be related to differential functions and/or protein-protein interactions of ATOX1 in myoblasts compared to cancer cells or species specific functions of ATOX1. Taken together, these data suggest that ATOX1 likely modulates myoblast proliferation by tuning copper availability via transport to ATP7A, though other ATOX1 functions may also contribute.

Indeed, detection of ATP7A as an ATOX1 proximal protein was most significant in myoblasts and myocytes, which is consistent with the importance of ATP7A in early differentiation that we previously reported in muscle and others have reported in neuronal cells.^10,12,13^ The ATOX1 proximal proteins identified in myocytes by comparative proteomics included multiple proteins related to endosomal trafficking, cytoskeleton, and post-Golgi sorting, all of which are consistent with ATOX1 functioning to deliver copper to ATP7A. In fact, several ATOX1 proximal proteins found in myoblasts and myocytes were also detected in proximity labeling and standard immunoprecipitation studies to probe the protein binding partners of ATP7A including COPA, SNX9, and SCAMP3.^41,42^ ATP7A itself was not detected since boiling during the streptavidin elution step renders the ATP7A protein insoluble. Although ATOX1 deficiency caused only a minor differentiation defect, cells plated at low density showed a stronger phenotype, which is consistent with the role of ATOX1 in delivering copper to ATP7A for assembly into secreted cuproproteins like lysyl oxidase (LOX).^12^ By contrast, cells plated at high density likely produce sufficient LOX and other cuproproteins to promote myotube formation. Thus, in intermediate stages of tissue differentiation, canonical ATOX1 functions likely contribute to copper delivery to tissue-specific and/or differentiation associated secreted cuproproteins via ATP7A.

Consistent with previous findings suggesting that ATOX1 can localize to the nucleus, several nucleic acid binding proteins were among the detected ATOX1 proximal binding partners.^24,43,44^ Our findings confirm results from a previous study identifying the transcription factor CRIP2 as an ATOX1 and copper binding protein in a study that employed APEX2 tagged ATOX1 with an added nuclear localization signal (NLS). CRIP2 has also been reported as a copper responsive transcription factor essential for myoblast differentiation.^31^ Thus, our data support a model wherein ATOX1, even in the absence of exogenous NLS, may deliver copper to CRIP2 and other transcription factors in myoblasts and myocytes. Interestingly, a number of RNA binding proteins were also detected as ATOX1 proximal proteins in this work in line with the previous study of NLS-tagged ATOX1-APEX2. Indeed, RNA binding proteins were particularly enriched in ATOX1 proximal proteins in myotubes. Although RNA binding proteins are often identified as contaminants in proteomics experiments, they tend to be expressed at much lower levels in fully differentiated myotubes compared to myoblasts and myocytes.^45,46,47,48^

Additional studies of ATOX1-RNA binding protein interactions are needed to better understand the functional relevance of copper and/or ATOX1 interactions with RNA binding proteins in skeletal muscle.

This study reveals shifting binding partners and differential functions of ATOX1 during myoblast differentiation. However, several limitations should be noted. Although proteins detected in minus biotin phenol negative controls were eliminated, it is probable that some APEX2-ATOX1 proximal proteins do not represent true ATOX1 binding partners. Additionally, the necessary use of hydrogen peroxide for proximal protein labeling by APEX2 may impair copper binding to ATOX1 and disrupt its protein-protein interactions. Nevertheless, the ATOX1- ATP7A interaction, which likely requires copper, is readily detectable using APEX2-ATOX1.^43,49^ Finally, given that ATOX1 is a small protein (∼7 kDa), fusion to the larger APEX2 protein could influence its localization, function, or complex formation. While detection of the expected ATOX1 localization and known binding partners should strengthen confidence in these results, further studies are needed to more precisely characterize stable and transient ATOX1 interactions with candidate partners.

## Materials and Methods

### Growth and differentiation of cell lines in culture

C2C12 immortalized skeletal myoblasts (purchased from American Type Culture Collection [ATCC]) and HEK293T cells (a kind gift from Dr. Bill Miller, University of Cincinnati) were grown in growth medium (GM) consisting of Dulbecco’s Modified Eagle Medium (DMEM, Corning, MT10013CV) in the presence of 10% fetal bovine serum (FBS, Cytiva SH3091003), 50 µg/ml penicillin and streptomycin (pen/strep, Corning MT30001CI), and 2.5 µg/ml Plasmocin treatment (InVivogen, NC9698402) at 37 °C in 5% CO2. C2C12 myoblasts were induced to differentiate by growing to about 90% confluency in GM then switching to differentiation media (DM) consisting of DMEM with 2% horse serum (Cytiva, SH3007403) and 50 µg/ml pen/strep and incubating at 37 °C in 5% CO2. Myocytes were harvested after 48hr of incubation while myotubes were harvest after 72-96 hours of incubation once formation of large myotubes was clearly visible. To generate myotubes from low density cells, C2C12 myoblasts were plated and allowed to grow to ∼50% confluence and induced to differentiate. For cell counting, equal numbers of cells were plated and counted 48 hours after inducing *Atox1* knockdown.

### Plasmids and cloning

The sequence encoding the APEX2-ATOX1 construct was synthesized by Azenta/Genewiz and subcloned from commercial vector into pcDNA 3.1 (ThermoFisher V79020) and then into the lentiviral vector pCW57.1 using EcoRI restriction enzyme recognition sites. The pCW57.1 was a gift from David Root (Addgene plasmid # 41393; http://n2t.net/addgene:41393; RRID:Addgene_41393). Plasmids were confirmed by Sanger sequencing and whole plasmid sequencing.

### Immunoblot analysis

All immunoblots were loaded with 10 µg of protein unless otherwise noted. Protein samples were separated on 4-20% TGX stain free protein gels (Bio-Rad, 4568094) and then transferred to nitrocellulose membrane (Cytiva, 45-004-001) using a Bio-Rad Transblot (1704150) following the manufacturer’s instructions. Membranes were blocked for 5-10 minutes in Everyblot blocking buffer (Bio-Rad 12010020) at room temperature with agitation. Blots were incubated with appropriate primary antibody for 1 hour at room temperature or overnight at 4°C with the (Supplemental Table 1). After primary antibody incubation, membranes were washed three times for 5 minutes with TBST (20mM Tris, 150mM NaCl, 1% Tween-20, pH 7.6) before incubation in appropriate secondary antibody for either 1 hour at room temperature or overnight at 4°C. Membranes were then washed three times for 5 minutes with TBST and developed with either Pico Plus ECL substrate (Thermo, PI34580) or Femto Plus ECL substrate (Thermo, PI34095) for 1 minute at room temperature before being visualized with a Bio-Rad Chemi-doc imager.

Relative protein expression was quantified by densitometry using Bio-Rad Image Lab software and statistical analysis was completed using GraphPad Prism.

### Generation of doxycycline-inducible APEX2-ATOX stable cell line

Six well plates were treated with Matrigel (Corning, 354234) diluted in DMEM for 1 hour at ambient temperature and washed with PBS. 3.5 x 10^5^ HEK 293T cells were seeded per well and incubated overnight. Approximately 16-18 hours after seeding, medium on HEK 293T cells was changed to growth medium without penicillin-streptomycin (transfection medium). Plasmids encoding APEX2-ATOX1 were co-transfected alongside PsPax2, a packaging construct containing HIV GAG-Pol, and PMDG.2, a viral envelope expressing plasmid containing VSVG envelope at a ratio of 4:2:1. After approximately 15-16 hours, transfection media was removed, and growth media was added to each well. To test for transfection efficiency, puromycin selection was used by adding puromycin (2 ug/ml final concentration) to each well.

Approximately 24 hours after addition of puromycin, low passage C2C12 myoblasts were seeded into 10 CM plates at low density. 48-72 hours later, enriched medium was removed from HEK293Ts and immediately placed on ice. Fresh growth medium was added back onto HEK293Ts to collect and store a second round of vector. Enriched medium was centrifuged twice at 500xg for 5 minutes to remove cellular debris, moving the supernatant to a fresh tube after the first spin, and further clarifying by filtration through a Whatman PES 0.45 µm syringe filter. In growth medium containing polybrene (8 ug/ml final) 3mLs of enriched medium containing lentiviral particles were added. Remaining vector was frozen overnight in a Styrofoam container at -80 C and moved to liquid nitrogen the following day. To ensure proper distribution of vector, plates that received enriched medium were rocked back and forth repeatedly every 30 minutes for eight hours. Approximately 24 hours after addition of APEX2- ATOX1 containing vector, fresh growth medium containing puromycin (2 ug/ml final) was added onto transduced C2C12 myoblasts and again 48 hours after transduction. Selection pressure was maintained by treating with puromycin for 48 hours after every passage or upon thawing fresh cell stocks.

### Induced expression of doxycycline inducible APEX2-ATOX1 and labeling

For labeling experiments in proliferating myoblasts, transduced cells were grown to about 60% confluency in GM. Doxycycline (Fisher, BP26535) was then added to a final concentration of 2.5 µg/mL (optimal concentration determined through titration Fig. 2) and incubated 24 hr. Cells were then incubated in GM with biotin-phenol (Sigma, SML2135-50MG) at a final concentration of 2.5mM for 30 min at 37 °C in 5% CO2. After incubation, cells were washed two times with sterile PBS then incubated in 0.75mM H2O2 (Fisher, H325-500) for exactly 1 minute. The reaction was quenched by washing cells two times with stop/wash buffer (PBS supplemented with 10mM MgCl2, 20mM CaCl2, 10mM NaN3, 5mM Trolox, 10 mM sodium ascorbate) followed by two more washes of sterile PBS before total cell lysates were collected in 250-500µL of RIPA lysis buffer for analysis. Labeling experiments in myocytes and myotubes followed the same protocol as described above with the following modifications. Transduced cells for myocyte experiments were grown to about 90% confluency in GM before being switched to DM for 24hr before doxycycline treatment. Transduced cells for myotube experiments were grown to 90% confluency in GM before switching to DM for 72-96hr before doxycycline treatment.

### Cell fractionation

Medium was aspirated from cells and then washed 1x with cold PBS. Cells were then scraped into 1 mL of cold PBS with a rubber scraper and centrifuged for 5 minutes at 500 x g. After centrifugation, supernatant was removed and 100 µL of NP-40 lysis buffer (50mM Tris HCl pH 7.4, 150 mM NaCl, 1% NP-40, 5mM EDTA) was added and mixed by gently pipetting up and down. Cells were then incubated for 5 min on ice with occasional gentle mixing. After incubation, 30 - 40 µL of total lysate was collected as the input fraction while the remaining cell lysate was centrifuged for 5 minutes at 500 x g. After centrifugation, the supernatant was collected as the cytosolic fraction. The remaining nuclear fraction pellet was washed with 100 µL of cold PBS by gently pipetting up and down and then centrifuged for 5 minutes at 500 x g. The supernatant was removed, and the nuclear pellet fraction was resuspended in 60 - 70 µL of RIPA lysis buffer. All samples were then either frozen at -80°C for long-term storage or sonicated on ice with an ultrasonic membrane disruptor (Fisher Model 100) 2 times for 5-10 seconds each and then centrifuged for 30 min at 21,000 x g in preparation for SDS-PAGE and immunoblotting.

### Streptavidin affinity pulldown and analysis

Total cell lysates were scraped into 250 µL of RIPA lysis buffer (25 mM Tris HCl pH 7.6, 150mM NaCl, 1% NP-40, 1% Sodium Deoxycholate, 0.1% SDS) supplemented with protease inhibitor (Pierce, PIA32953) and incubated on ice for 30 min. Cell lysates were sonicated on ice with an ultrasonic membrane disruptor (Fisher Model 100) 2 times for 5-10 seconds each and centrifuged at 21,000 x g for 30 minutes. Supernatants were transferred to new tubes and protein concentration was determined through Bradford assay according to the manufacturer’s instructions (Bio-Rad, 5000205). 100 µL of Pierce high-capacity streptavidin agarose beads (PI20357) per sample were washed twice with 500µL of IP buffer (50mM Tris HCl pH 7.5, 150 mM NaCl, 5mM MgCl2, 1% NP-40) supplemented with protease inhibitor (Pierce) with rotation for 5 minutes. Beads were pelleted by centrifugation for 1 min at 500 x g and blocked for 2-3 hours in 500µL of 1x IP blocking buffer (IP Buffer + 1% Bovine Serum Albumin (Fisher: BP9703-100) at 4°C. Beads were pelleted by centrifugation and then washed 2 x 5 minutes in IP Buffer. 500µg of protein lysates were added to agarose beads and incubated overnight with rotation at 4°C. After incubation, bead slurry was washed 5x for 5 minutes with 500µL of IP buffer with rotation. Proteins were eluted from beads by adding 50 µL of 2X Laemmli sample buffer (Bio-Rad, 161-0747) and 50 µL of IP buffer supplemented with 6.5 M urea (Fisher, 434720010) followed by incubation at 95°C for 5 minutes. Total protein recovered was evaluated via silver stain using 4-20% TGX stain free gels (Bio-Rad, 4568094). After separation, protein samples were stained following the manufacturer’s instruction (Thermo-Fisher, 24612) and visualized with a Bio-Rad Chemi-doc imager. Biotinylation efficiency was determined via immunoblot analysis using streptavidin – HRP (Supplemental Table 1).

### LC-MSMS for ATOX 1 proximity profiling and functional annotation

Streptavidin eluates from APEX2-ATOX1 expressing myoblasts, myocytes and myotubes from three independent experiments were used for proteomic profiling. Cells without biotin phenol treatment were used as negative controls. Enriched biotinylated proteins in Laemmli sample buffer were prepared for and subjected to in-gel trypsin direction and recovery followed by a label-free quantitative nanoLC-MS/MS workflow as detailed previously (Bennett). Briefly samples were all run about 2 cm into an Invitrogen 4-12% Bis-Tris gel using MOPS buffer with pre-stained molecular weight marker lanes in between. The full 2 cm protein sections were excised, reduced with dithiothreitol, alkylated with iodoacetic acid, and digested overnight with trypsin. The peptides were subsequently recovered followed by nanoLC-MSMS on a Dionex Ultimate 3000 RSLCnano coupled to a Thermo Orbitrap Eclipse mass spectrometry system. Data were collected using Xcaliber 4.3 software (ThermoScientific) with label free quantitation comparative profiling of proteins detected from each sample group achieved using Proteome Discoverer 3.0 (ThermoScientific) against the complete *Mus musculus* protein database. Proteins with the minimum of 2 high (99%) confidence peptides with significant proteins differences (p<0.05) and minimum of 2-fold change between the groups are reported. Proteins detected in all three replicates and not detected in -BP controls were used for downstream functional annotation using DAVID and STRING databases.

### Crosslinking Reverse Co-Immunoprecipitation

C2C12 cells were grown to about 80% then washed with 1x PBS twice and crosslinked with 0.1mM DSP (ThermoScientific 22585) in PBS for 30 min at 37°C. Cells were rinsed twice with PBS and the reaction quenched with 20 mM Tris-HCl (VWR 1185-53-1) for 15 min at room temperature. Cells were then rinsed twice more with 1x PBS and scraped into 500 µL of RIPA lysis buffer for total cell lysate preparation as described below. Total cell lysate protein concentration was calculated using Bio-Rad Bradford assay kit (Bio-Rad, 5000205). Protein A agarose beads (Pierce 22810) were used for immunoprecipitation of MEK1/2 (Supplemental Table 1). 35µL of beads were rinsed twice with IP buffer and then incubated with MEK1/2 antibody according to manufacturer recommendations overnight with rotation at 4°C. Protein lysates (250 – 500µg) were precleared in 25µL of rinsed beads for 1 hour at 4°C. After preclearing, cell lysates were incubated with bead-antibody mixtures overnight at 4°C with rotation. Beads were washed with 500 µL of IP buffer 5 x for 5 min each then incubated with 50µL of 2x Laemmli, 45µL IP buffer, and 5µL of 1M dithiothreitol (DTT) for 30 min at 37°C with periodic shaking. Bead mixture was then boiled for 5 min and then eluates were removed for analysis via immunoblot according to the protocol described below.

### Transfection and siRNA knockdowns

Knockdown of *Atox1* was performed a previously described.^12^ Briefly, dicer substrate siRNAs targeting the *Atox1* 3’ UTR were purchased from IDT. Myoblasts growing in transfection medium (growth medium without penicillin-streptomycin) were transfected with *Atox1* targeting siRNAs or non-targeting negative control siRNAs using Lipofectamine 3000 (ThermoFisher L3000008). Transfecting cells were refreshed with fresh growth medium or differentiation medium without transfection mix after ∼16 hours in transfection mix. Knockdown of ATOX1 was confirmed by immunoblot.

### Quantitative reverse transcriptase PCR (qRT-PCR)

RNA was isolated using TRIzol (Invitrogen 15596018) according to manufacturer’s instructions. cDNA was synthesized using the Maxima First Strand cDNA Synthesis kit (Fisher K1672) according to manufacturer’s instructions. Quantitative reverse transcriptase PCR was then performed using SYBR Select Master Mix (Applied biosystems 4472919) on a QuantStudio 3 Real-Time PCR System (Applied Biosystems) using primers targeting *Atox1*(F: *cgagttctccgtggacatga* R: *cctcccagcttgttgaggac*) and *Rplp0* (F: *gggcgacctggaagtccaact* R: *cccatcagcaccacagccttc*). Results were quantified using comparative Ct method with *Rplp0* as a normalizer.^50^

### EZ-link maleimide labeling

Maleimide labeling of free cysteines in ATOX1 was performed as previously described.^10^ Cells were grown to desired stage of differentiation then scraped into MOPS lysis buffer (50 mM MOPS pH 7, 500 mM sucrose, 300 mM NaCl) and incubated on ice for 10 min. Cells were then sonicated and protein concentration was determined as described above. Oxidized sample was incubated in 1 mM H2O2 for 5 min at room temperature before labeling. Samples were labeled by adding 1 µL of 250 mM EZ-link Maleimide (Thermo 21911) stock reagent to 99 µL of protein lysates for a final concentration of 2.5 mM. Mixtures were incubated on ice for 3 hours and then quenched by addition of 10 µL of 1 M L – Cysteine. 10 -15 µg of labeled protein lysates were analysis via immunoblot.

### Immunofluorescence staining

Cells were washed two times with 1x Phosphate-buffered saline (PBS: 137 mM NaCl, 2.7 mM KCl, 10 mM Na2HPO4, 1.8 mM KH2PO4, Fisher: BP665-1) and fixed with 3.7% formaldehyde (Fisher, BP531500) for 10 min at room temperature. Cells were washed twice with PBS then permeabilized in PBS supplemented with 0.3% Triton-X (Fisher, BP151 100) by incubating at room temperature for 15 minutes. After incubation, cells were rinsed once with PBS and incubated with IF Blocking buffer (PBS supplemented with 0.1% Triton-X and 3% bovine serum albumin [Fisher, BP9703-100]) for 1 - 2 hours at 4°C. After blocking, cells were rinsed once with PBS then incubated with primary antibody diluted in 0.5X IF blocking buffer overnight at 4°C (antibody information can be found in Supplemental Table 1). After incubation cells were washed three times with 0.5X IF blocking buffer then incubated with secondary antibody diluted in 0.5X IF blocking buffer for 1-2 hours at room temperature protected from light. After incubation cells were washed three times with 0.5X IF blocking buffer then incubated with 4’,6- diamidino-2-phenylindole (DAPI: Fisher, D3571) diluted 1:1000 in PBS for 5 - 10 minutes. After incubation, cells were washed twice with PBS and kept in fresh PBS while imaging. Cells were imaged with an Olympus IX83 inverted fluorescence microscope using a U Plan fluorite 10x objective lens (NA 0.3 WD 10 mm). Images were captured using a DP74 Color CMOS Camera (cooled 20.8 MP pixel-shift, 60 FPS) using CellSens Dimension V2 software. Images were subjected to 2D-deconvolution and exported as red, green, blue (RGB) TIF files for analysis in FIJI. Three images were taken at random positions for each sample and experiments were completed in triplicate.

### Quantification of differentiation

Fully formed myotubes were stained with anti-eMyHC and DAPI. Three images were taken at random positions in the well for each sample and exported as red, green, blue tiff images to FIJI. Fusion index was determined by calculating the ratio of nuclei in eMyHC positive cells to total nuclei in each image. Fusion index calculations from each image were averaged and analyzed by ANOVA in GraphPad Prism version 10.4.0. Myotube width was measured in FIJI by choosing 10 myotubes randomly for each image and quantifying width in nm. Fusion index and myotube width analyses were performed blinded.

## Acknowledgements

The authors would like to acknowledge the University of Cincinnati core facilities employed to generate data reported here. Mass spectrometry data were collected and analyzed in the University of Cincinnati College of Medicine Proteomics Laboratory under the direction of KD Greis, Ph.D. This work was supported by the University of Cincinnati Research Council Research Scholars Award to K.E.V. and National Institutes of Health (NIH) 5R35GM146878 to K.E.V. and 5R01CA262664 and 5R01DK131190 to M.J.P.

## Data Availability Statement

Proteomics data used for this study openly available on figshare https://doi.org/10.6084/m9.figshare.29497352.v1

---

**Supplemental Figure 1:**
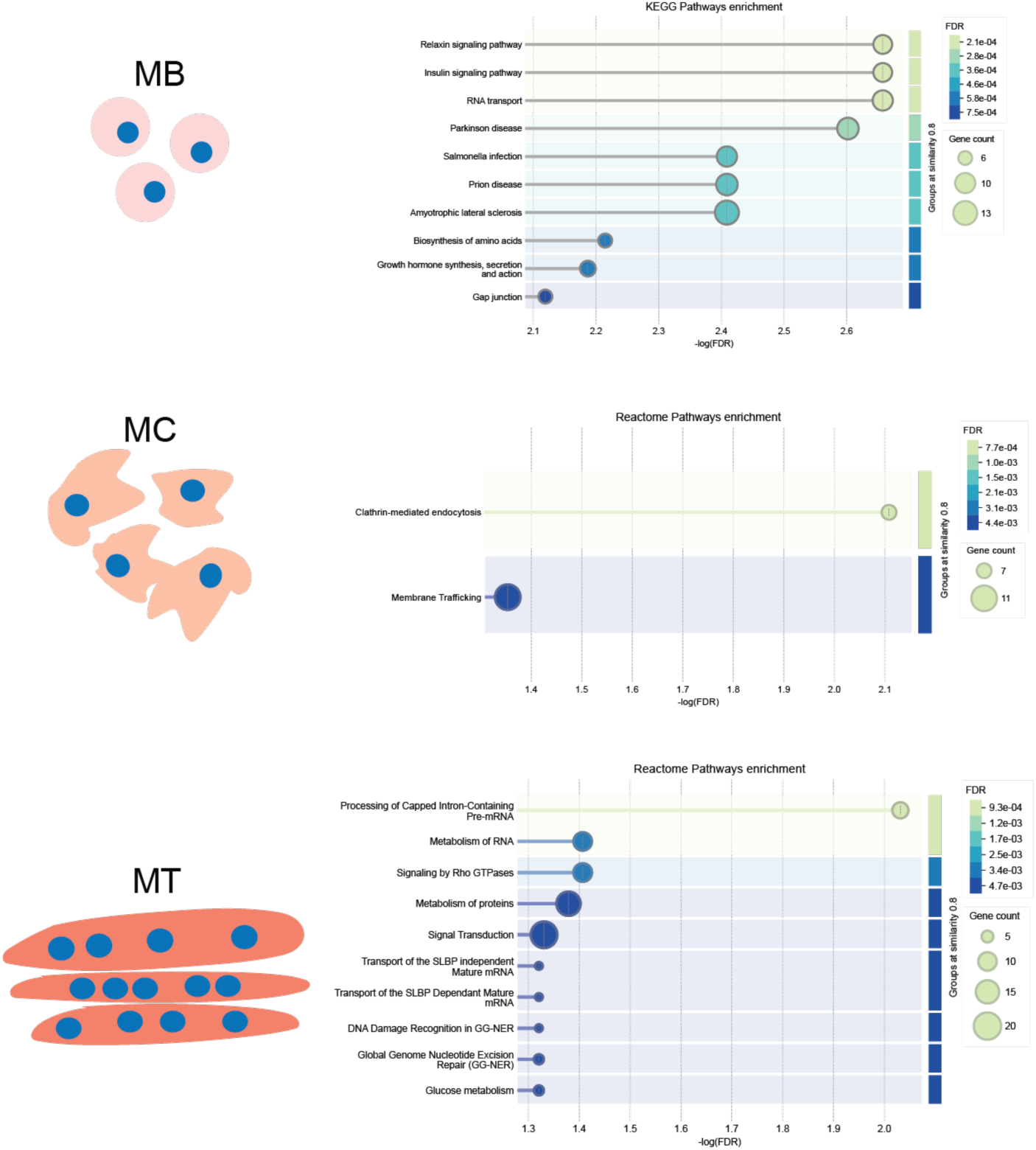
Additional functional annotation of ATOX1 proximal proteins. Shown are enriched KEGG pathway in myoblasts (MB) and Reactome pathways in myocytes (MC) and myotubes (MT). No KEGG pathway enrichment was detected in MC or MT and no Reactome pathway enrichment was detected in MB. Pathway analysis was performed using the STRING database.

**Supplemental Figure 2:**
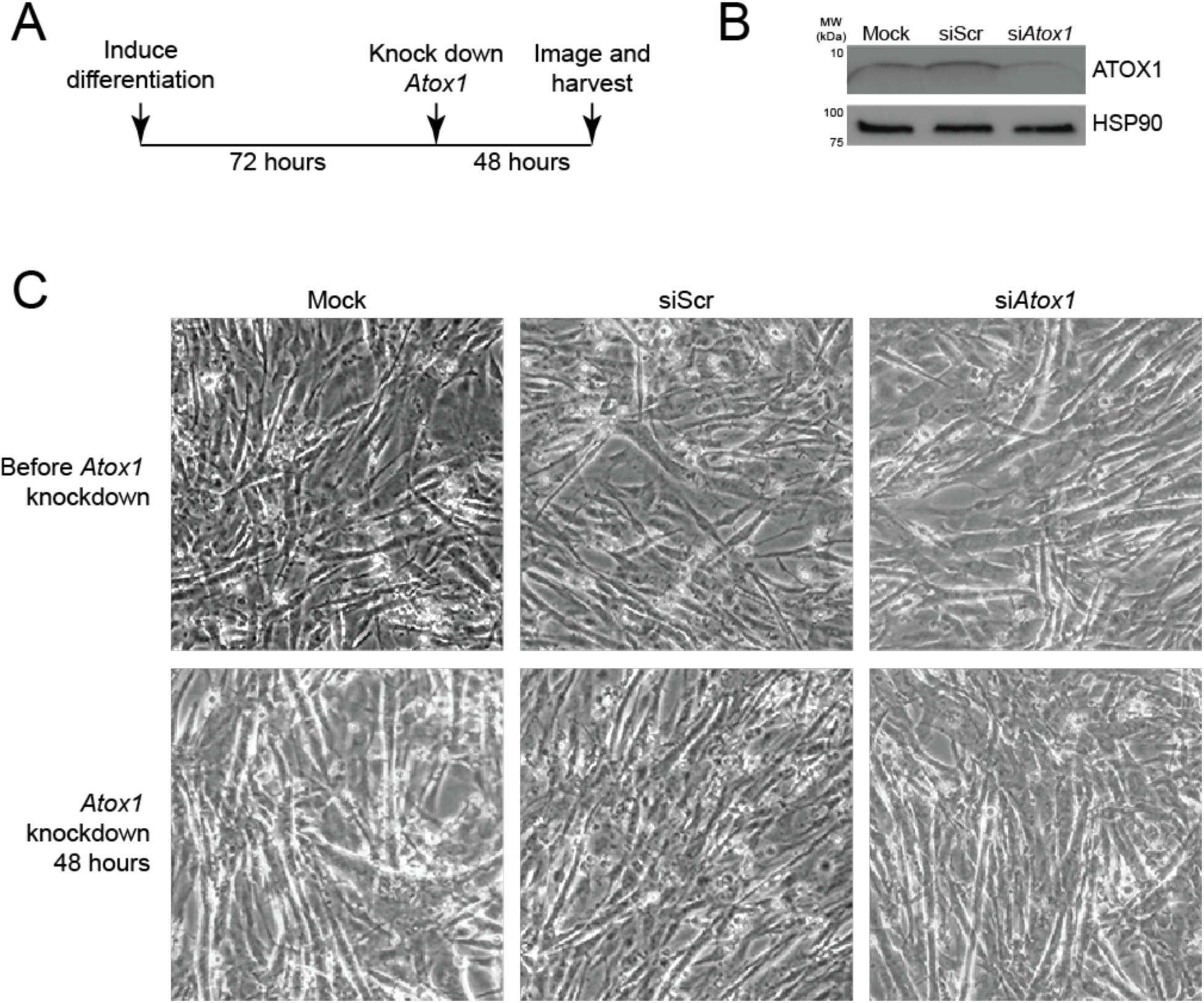
**No overt phenotype cause by ATOX1 deficiency in fully differentiated myotubes. *A)*** Schematic of experiment. Briefly, myoblasts were allowed to differentiate for 72 hours, transfected with control or *Atox1* targeting siRNA (siAtox1), and harvested 48 hours later. ***B)*** Immunoblot probed with an antibody to ATOX1 showing reduced ATOX1 protein in *Atox1* knockdown myotubes compared to mock transfected (Mock) or non-targeting siRNA (siScr) control myotubes. Antibody targeting HSP90 was used as a loading control. ***C)*** Phase contrast images of early (top) and mature (bottom) myotubes in cells prior to transfection (top) and 48 hours after transfection (bottom) showing no overt phenotype in myotubes transfected with siAtox1.

**Supplemental Table 1.**

| Company | Antibody | Catalog # | Dilution |
| --- | --- | --- | --- |
| Wang et al PLOS One, 2012 | ATP7A | N/A | 1:1000 |
| Proteintech | ATOX1 | Catalog#: 22641-1-AP | 1:1000 |
| Lab made | eMyHC | N/A | 1:10 |
| Lab made | Flag | N/A | 1:1000 |
| Cell Signaling Technology | Histone H3 | 4499s | 1:4000 |
| Cell Signaling Technology | GAPDH | 5174S | 1:1000 |
| Santa Cruz | CCS | sc-55561 | 1:1000 |
| Cell Signaling Technology | MEK1/2 | 9122 | 1:1000 |
| Lab made | ATP7A | N/A | 1:1000 |
| Cell Signaling Technology | HSP90 | 4877S | 1:4000 |
| Proteintech | PLOD2 | 21214-1-AP | 1:1000 |
| Proteintech | Scamp3 | 67932-1-Ig | 1:1000 |
| Proteintech | CRIP2 | 14801-1-AP | 1:1000 |
| Proteintech | SYNCRIP | 14024-1-AP | 1:1000 |
| Jackson Laboratory | Mouse-HRP | 115-035-003 | 1:10000 |
| Jackson Laboratory | Rabbit-HRP | 111-035-003 | 1:10000 |
| Cell Signaling Technology | Streptavidin-HRP | 3999S |  |
| Jackson Laboratory | Mouse-Alexa Fluor 488 | 715-545-151 | 1:500 |

